# Correspondence between population coding and psychophysical scaling models of working memory

**DOI:** 10.1101/699884

**Authors:** Paul M Bays

**Affiliations:** University of Cambridge, Department of Psychology, Cambridge, CB2 3EB, UK

## Abstract

A mathematical idealization of the way neural populations encode sensory information has been found to provide a parsimonious account of errors made by human observers on perceptual and short-term memory tasks. This includes the effects of set size and flexible prioritization of items within a set (Bays, 2014), the frequency and identity of “swap” or misbinding errors (Schneegans & Bays, 2017), subjective judgments of confidence (Bays, 2016; van den Berg et al., 2017), and biases and variation in precision due to serial dependent and stimulus-specific effects (Bliss et al., 2017; Taylor & Bays, 2018). A superficially quite different account of short-term recall has recently been proposed in work by Schurgin et al. (2018), who argue that taking into account the differences between physical and perceptual distance in a feature space reduces recall to a classical signal detection problem. Here I document a remarkable similarity between the two models, demonstrating a favourable convergence of neural- and cognitive-level accounts of working memory.

---

Information from the sensory environment is encoded in the brain in the activity of neural populations. Computational theories of population coding (Dayan & Abbott, 2001; Pouget et al., 2000) provide a link between single cell responses to stimuli observed in electrophysiological studies and the behavioral responses studied with psychophysical experiments. In many cases features of population codes have direct analogues in ideal observer and signal detection theories (Ma, 2010).

In a general population coding framework, stimulus features ***θ*** are encoded in the joint activity, **r**, of a set of *M* idealized neurons with responses determined by each neuron’s individual tuning in the feature space, 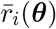, plus zero-mean random noise ***ε***,

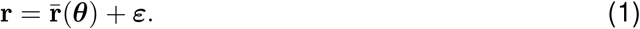

An estimate of the encoded feature values can be recovered from the population responses by finding the features *θ*′ that maximize the likelihood function,

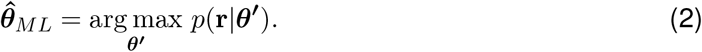

In previous studies (e.g. Bays, 2014; Schneegans & Bays, 2017; Taylor & Bays, 2018) it has been shown that versions of this model provide a good account of errors observed in recall of elementary visual features (e.g. stimulus orientations), and in particular the effect on those errors of manipulating set size, i.e. the number *N* of features to be stored in memory. The specific model proposed in Bays (2014) assumed each stimulus feature, *θ* ∈ [−*π*, *π*) was encoded by an independent population of *M* neurons with identical tuning functions, *f* (*x*), scaled by an amplitude *ξ*, and translated such that the preferred feature values of the neurons, *φ*_*i*_, were uniformly distributed throughout the circular space of possible features, such that the mean response of the *i*th neuron was,

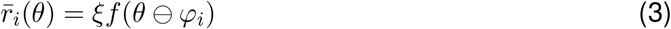

where ϴ indicates subtraction on the circle, and the total expected response amplitude of all neurons encoding all stimuli was normalized, i.e. 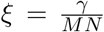, where *γ* is a population gain parameter. A von Mises tuning function was used in the main part of the modelling, *f* (*x*) = exp(*κ*(cos(*x*) − 1)) (with *κ* a free parameter), although the model predictions do not depend strongly on the shape of this function (Bays, 2014).

In Bays (2014), responses *r*_*i*_ were corrupted by independent Poisson noise, however very similar results have been obtained using a Gaussian approximation to Poisson (e.g. Schneegans & Bays, 2017), i.e. 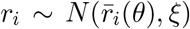. In this case, and assuming there are sufficiently many neurons to densely cover the space, the ML decoder is given by

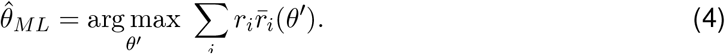

## Psychophysical scaling model

According to the psychophysical scaling account of visual memory errors proposed by Schurgin et al. (2018), the distribution of recalled feature values depends on the perceptual distance between points in the stimulus space, characterized by a non-linear increasing function of distance, *g*(·), that can be estimated from a purely perceptual task. Given a stimulus feature *θ*, a noisy “memory match” signal, 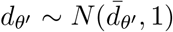, is generated for each possible response *θ*′ as a draw from a unit-variance normal distribution with mean,

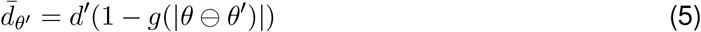

where *d*′ is a free parameter of the model. The response is then chosen as the feature value with the highest signal strength (i.e. a “winner-takes-all” decision rule),

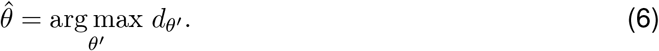

To demonstrate the close correspondence between this account and the population coding model described above, we need only define a symmetrical function *f* (*x*) = 1 − *g*(|*x*|). Then,

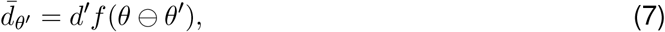

and it can be seen that the mean memory match signals 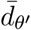, in Eq 7 are identical to the mean neural responses 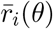, in Eq 3, with the possible responses *θ*′ corresponding to the preferred stimulus values *φ*_*i*_, and *d*′ equal to the peak response amplitude *ξ*.

The psychophysical scaling model assumes noise with unit variance, 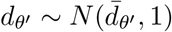. However the neural responses in the population coding model can be scaled arbitrarily without changing the predicted errors, as long as the ratio of signal to noise remains unchanged. If we scale by a factor 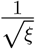, then 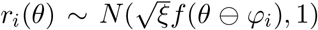, and we can see that setting 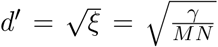 makes the distributions of signal strengths in the two models identical as well.

## Synthesizing the models

Although conceptualized in terms of signal detection theory, the psychophysical scaling model has a plausible biological foundation in computational theories of neural population coding. According to the neural formulation, the perceptual similarity between features relates directly to the tuning functions in the underlying populations responsive to those features. The variable “memory match” signal indicating the strength of internal evidence for a particular feature value has a neural counterpart in the noisy activity of neurons tuned to prefer that same feature value.

The only significant discrepancies between the model proposed by Schurgin et al. (2018) and the model of Bays (2014) – which has been shown previously to quantitatively account for many aspects of human recall performance – is that the former model uses a different (and in general sub-optimal) decision rule (winner-takes-all, Eq 6, instead of maximum likelihood, Eq 4) and at this point does not specify how *d*′ varies with set size (the population coding model predicts *d*′ proportional to 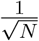, which we note is also consistent with predictions of the sample-size model of Shaw, 1980, indicative of the deep theoretical connections between these models).

For the correspondence between models to be exact, the perceptual distance function in the psychophysical scaling model, *g*(*x*), which Schurgin et al. (2018) estimated non-parametrically from the results of a separate perceptual task, should equal one minus the tuning function *f* (*x*) in the neural model (for which a von Mises function has been specified). However, from a population coding perspective, a more natural choice to describe perceptual distance between two features would be based on the differences in the mean neural responses they elicit; one concrete possibility is: 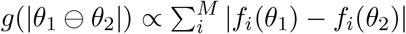.

I predict that incorporating an appropriate function of this kind into the population coding model may allow it to simultaneously fit data from both perceptual scaling and working memory tasks. This could potentially provide a mechanistic basis for a perceptual distance function having the particular form that is observed empirically, something which the psychophysical scaling model does not currently provide. In particular, a key observation of Schurgin et al. (2018), that participants fail to discern in a perceptual task whether an item 120° or 180° from a target in color space is closer to the target, could reflect the fact that the neurons in visual areas that are active in response to the colors at 120° or 180° have negligible overlap with the neurons that respond to the target color at 0°.

## Conclusion

As famously recognised by Marr (1982), the challenge of understanding a system as complex as the brain is unlikely to be met by focusing on just one level of analysis or explanation. In this case, the psychophysical scaling account may be viewed as providing a new cognitive or algorithmic-level description of the same short-term memory mechanisms specified previously at a biophysical or implementation level by the population coding model. This synthesis may lead to fruitful directions for future research.

## Acknowledgments

I thank Sebastian Schneegans for comments on the manuscript. This work was supported by the Wellcome Trust (Grant number 106926).

